# Polycomb Repressive Complex 2-controlled Essrg regulates intestinal Microfold cell differentiation

**DOI:** 10.1101/2020.11.13.379610

**Authors:** Joel Johnson George, Mikko Oittinen, Laura Martin-Diaz, Veronika Zapilko, Sharif Iqbal, Terhi Rintakangas, Fábio Tadeu Arrojo Martins, Henri Niskanen, Pekka Katajisto, Minna Kaikkonen, Keijo Viiri

## Abstract

Microfold cells (M cells) are immunosurveillance epithelial cells located in the Peyer’s patches in the intestine responsible for monitoring and transcytosis of antigens, microorganisms and pathogens. Many transcription factors, e.g., Spi-B and Sox8, necessary to M cell differentiation have been described but the exhaustive set of factors sufficient for differentiation and development of a mature M cell remains elusive. Moreover, the role of polycomb repressive complex 2 (PRC2) as an epigenetic regulator of M cell development has not yet been interrogated. Here, we show that PRC2 regulates a significant set of genes during the M cell differentiation including many transcription factors. Estrogen related receptor gamma (Esrrg) is a novel M cell specific transcription factor acting on a RankL-Rank induced NF-kB pathway, upstream of Sox8 and necessary but not sufficient for a mature M cell marker Gp2 expression. To conclude, with the aid of PRC2 target survey we identified the list of developmental genes specifically implicated in M cell development and Essrg as a necessary factor for Sox8-mediated M cell differentiation.

## Introduction

The Gut-associated lymphoid tissue (GALT) is involved in immune surveillance of antigens, microorganisms and foreign pathogens that constantly thrive in the mucosal surface of the intestinal tract. The GALT is the immune initiation site against mucosal antigens and houses specialized gut immune epithelial cells known as M cells or Microfold cells, these cells envelop the luminal surface of GALT also known Peyer’s patches (PPs). (Mabbott et al., 2013; Neutra et al., 1996; Owen, 1999). The principal role of the M cells is the uptake and transcytosis of luminal antigens into the GALT as they have a high phagocytic and transcytosis capacity, which is responsible for the rapid transport of bacterial antigens to antigen-presenting immature dendritic cells (Neutra et al., 2001; Rios et al., 2016). Subsequently, these dendritic cells undergo maturation and activate antigen specific naive T cells, that support B cell activation, resulting ultimately in the generation of IgA-producing plasma cells (Kraehenbuhl & Neutra, 2000). It has been shown previously that the absence of M cells or their antigen uptake receptor glycoprotein 2 (Gp2) mitigates the mucosal immune responses by T cells in mice infected with *Salmonella enterica* serovar Typhimurium. This is predominantly due to lack of bacterial transcytosis by the mature GP2 receptor into the GALT (Kanaya et al., 2012; Kimura et al., 2019). Correspondingly, perturbances in transcytosis of *Yersinia enterocolitica* in PPs were observed in Aif1 mutant mice (Kishikawa et al., 2017). Recently, it was shown that M cells self-regulate their differentiation by expressing osteoprotegerin (OPG), a soluble inhibitor of RANKL, which suppresses the differentiation of adjacent FAE cells into M cells. This self-regulatory machinery of M cell density is necessary as *Opg*^−/−^ mice were highly susceptible to mucosal infection by pathogenic bacteria because of the augmentation of bacterial translocation via M cells (Kimura et al., 2020). Overall, defects in M cell-dependent antigen uptake led to a decrease in production of antigen-specific secretory IgA (S-IgA) in the gut (Hase et al., 2009; Kishikawa et al., 2017; Rios et al., 2016).

M cell differentiation of cycling intestinal crypt cells that express Lgr5 and RANK receptors is induced by the Receptor activator of nuclear factor-κB ligand (RankL). RankL is secreted by stromal cells also known as M-cell inducer cells (MCi cells) or immune cells in the sub epithelial dome (SED) (Mabbott et al., 2013; Nagashima et al., 2017). RankL deficient mice have very few M cells but exogenous administration of recombinant RankL was able to mitigate that loss (Knoop et al., 2009). RankL binding to Rank receptor leads to the activation of the intracellular adaptor molecule of RANK; TRAF6 which in turn leads to activation of both canonical (RelA/p50 heterodimer) and non-canonical NF-κB (RelB/p52 heterodimer) activation (Lernbecher et al., 1994; Walsh et al., 2014, Kanaya et al., 2018). The canonical RelA/p50 activation led to expression of early M cell markers like MarcksL1 and CCL9 whereas non-canonical RelB/p52 activation led to expression of Spi-B and Sox8 transcription factors, both deemed essential to maturation of M cells (Kanaya et al., 2018; Kimura et al., 2019). Along with RankL, expression of Spi-B and Sox8 are essential for the development of Gp2 positive M cells. Both Spi-B and Sox8 mutant mice exhibited the absence of mature M cells with Gp2 whereas Marcksl1^+^AnnexinV^+^ immature M cells were intact (de Lau et al., 2012; Kanaya et al., 2012; Kimura et al., 2019; Sato et al., 2013). Spi-B -/- still showed Sox8 expression and Sox8 -/- mice expressed Spi-B and even though both mice had activation of both NF-κB transcription pathway, p50/RelA and p52/RelB, they still showed an immature M cell phenotype lacking the expression of Gp2 (Kanaya et al., 2018). Taken together, despite their critical role in the onset of mucosal immune responses, M cell development and their differentiation into maturity have not yet been fully characterized partly due to their rarity in the gastrointestinal tract (Kanaya & Ohno., 2014). Importantly, the sole overexpression of Spi-B and Sox8 are not sufficient for the induction of GP2 receptor i.e., M cell maturation, suggesting that additional M cell specific transcription factors are needed (Kimura et al., 2019).

Intestinal cell differentiation, development and functionality are regulated by several factors, one of the indispensable ones being Polycomb Group (PcG) proteins. PcG proteins are essential for embryonic stem cell self-renewal and pluripotency but they are also necessary for the maintenance of cell identity and cell differentiation throughout life (Schuettengruber & Cavalli, 2009). They broadly form three groups of polycomb repressive complexes (PRCs) known as PRC1, PRC2 and Polycomb Repressive DeUBiquitinase, each of these complexes reassemble chromatin by explicitly defined mechanisms that involve variable configurations of core and accessory subunits. This configuration is demonstrated by the way PRC2 catalyzes trimethylation of histone H3 lysine 27 (H3K27me3) and presents a binding site for PRC1 in embryonic stem cells (Cao et al., 2002). Lee et al showed that PRC2 played a repressive role of expression of developmental regulators necessary for cell differentiation (Lee et al., 2006). Interestingly, genes critical for cell identity lose their methylation on H3 lysine K27 whereas genes that regulate alternate cell types keep their methylation and remain repressed (Bernstein et al., 2006). In human embryonic stem (ES) cells, Wnt-signaling genes are bound by PRC2, analogously this is also exhibited in adult tissues for instance in adipogenesis (Wang et al., 2010). The integrity and homeostasis of healthy intestine is partly regulated by canonical Wnt signaling and it has also been shown that secretory and absorptive progenitor cells show comparable levels of histone modifications at most of the same cis elements in the genome (T. H. Kim et al., 2014). Our study and others (Chiacchiera et al., 2016; Oittinen et al., 2017; Vizán et al., 2016) found out how PRC2 regulates a substantial subset of genes that were involved in canonical Wnt signaling and contributed to the differentiation of Lgr5 expressing stem cells to secretory and absorptive cell types in the intestine.

To further understand the complexity of M cell differentiation, we asked if PRC2 regulates M cell differentiation. We used high-throughput tools such as Chip-seq and Gro-seq to identify factors that contribute to the function and development of M cells in the intestine. We identified a total of 12 TFs that are regulated by PRC2 exclusively during M cell differentiation of which 6 were downregulated and 6 were upregulated. One of the M cell specific TF, Esrrg, was found to be essential for the differentiation of mature Gp2+ M cells in vitro. We found that Esrrg was expressed exclusively in M cells in Peyer's patches and was shown critical for the activation of Sox8 transcription factor. Esrrg expression was intact even in Sox8 deficient mice and was dependent on the activation of non-canonical NF-κB signaling. These observations illustrate that Esrrg is a crucial player in the differentiation and functionality of a mature Gp2+ M cell.

## Results

### PRC2 is not restricting M cell differentiation

We and others have previously shown that disrupting PRC2 activity leads to a precocious expression of terminal differentiation markers of intestinal enterocytes (Benoit et al., 2013; Oittinen et al., 2017; Vizán et al., 2016). Moreover, PRC2 has been shown to preserve intestinal progenitors and restricting also secretory cell differentiation (Chiacchiera et al., 2016). Contrary to absorptive cell differentiation (ENRI), when organoids were treated with RANKL the level of expression of PRC2 members Ezh2 and Suz12 were comparable to the levels in organoids grown in standard organoid culture media (ENR500) (Fig. 1A). Next, we asked if PRC2 inhibition (Fig. 1B) can augment M cell differentiation and we saw that, contrary to enterocyte differentiation, expression of all M cell markers are downregulated when the activity of PRC2 is pharmacologically inhibited during the RANKL-induced M cell differentiation (Fig. 1C).

**Figure 1.**
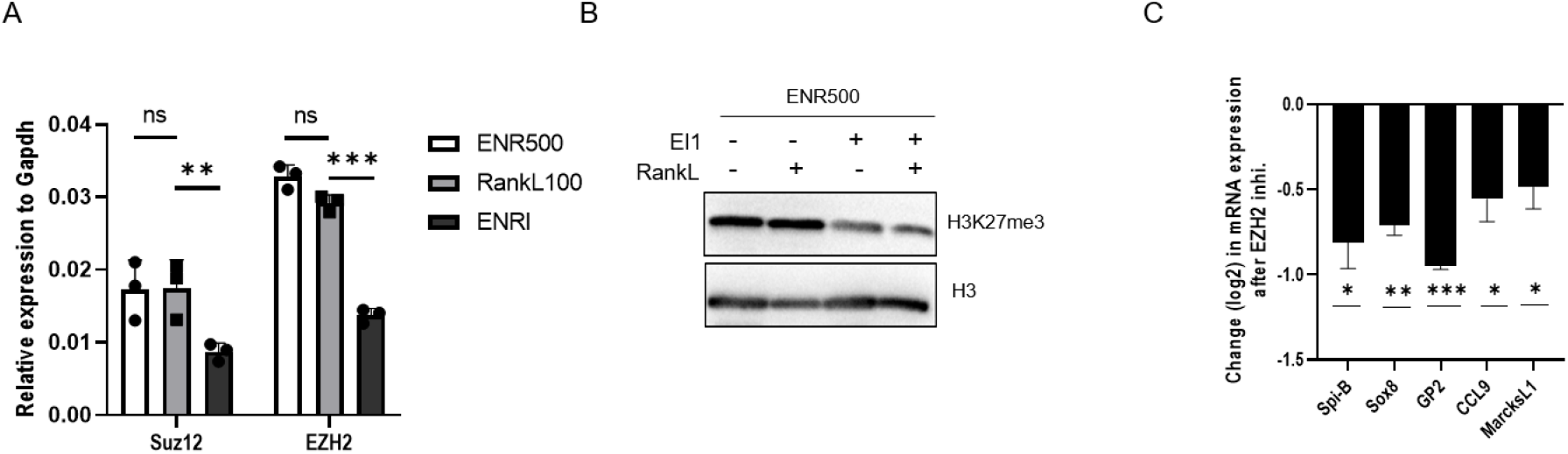
PRC2 members are expressed in M cells. A) RT-qPCR analyses of the expression of Suz12 and Ezh2 in mouse intestinal organoids grown in ENR, ENR+RankL and ENRI conditions. B) Immunoblot of H3K27me3 and H3 in organoids treated with EI1 inhibitor. Data are representative of 2 independent experiments. C) RT-qPCR analyses of the expression of M cell marker genes in RankL-treated organoids with or without EZH2 inhibitor EI1. Unpaired two-tailed Student’s *t* test was performed, n.s., not significant; *, P< 0.05; **; P < 0.01; ***, P < 0.005, *n* = 3. Values are presented as the mean ± SD

### PRC2 regulated genes during M cell differentiation

It has been previously shown that PRC2 regulates transcription factors necessary to intestinal stem cell maintenance and differentiation e.g., Ascl2 (Schuijers et al. 2015 & Oittinen et al. 2017) and Atoh1 (Chiacchiera et al. 2016). Since PRC2 mainly regulates genes involved in development or signaling (Ram et al. 2011 Cell) we reasoned that identifying genes regulated by PRC2 during M cell differentiation might reveal gene network necessary to this cell type in the intestine. M cell differentiation was induced in mouse intestinal organoids with RankL and gene expression was analysed with Gro-Seq and PRC2 target genes identified with ChIP-seq by using H3K27me3 antibody. Genes differentially expressed after RankL treatment are shown in figure 2A. ChIP-seq performed with H3K27me3 antibody revealed a significant number of genes regulated by PRC2 for M cell differentiation. When comparing our 3 different Chip-seq’s (WENRC – stem cell conditions; ENRI – enterocyte condition; RankL – M cell differentiation), we observed that in M cells, 38 (9.2%) and 35 (10.3%) of genes were upregulated but silenced by PRC2 in WENRC and ENRI, respectively. 32 PRC2-target genes were uniquely upregulated during M cell differentiation and not in enterocyte differentiation. 46 (27.7%) and 52 (11.5%) PRC2-target genes were silenced in organoids treated with RANKL but expressed in WENRC and ENRI, respectively. 42 genes were uniquely silenced in M cell but not in enterocyte differentiation. PRC2 target genes are shown in figure 2 B-D and the M cell specific accumulation and ablation of H3K27me3 signal in the gene promoters are shown in figure 2E. Gene ontology analyses indicate that PRC2 regulate many DNA-binding transcription factors during the M cell differentiation (Fig. 2F). Total of 12 transcription factors were differentially expressed: six were silenced by PRC2 in M cells and six were specifically expressed in M cells but repressed by PRC2 in organoids grown both in stemness and enterocyte conditions. The six PRC2-target genes specifically expressed in M cells are Sox8, Atoh8, Esrrg, Smad6, Maf and Zfp819. The six genes silenced are Hoxb5, Hoxb9, Sp9, Sp5, Nr4a1 and Atf3.

**Figure 2.**
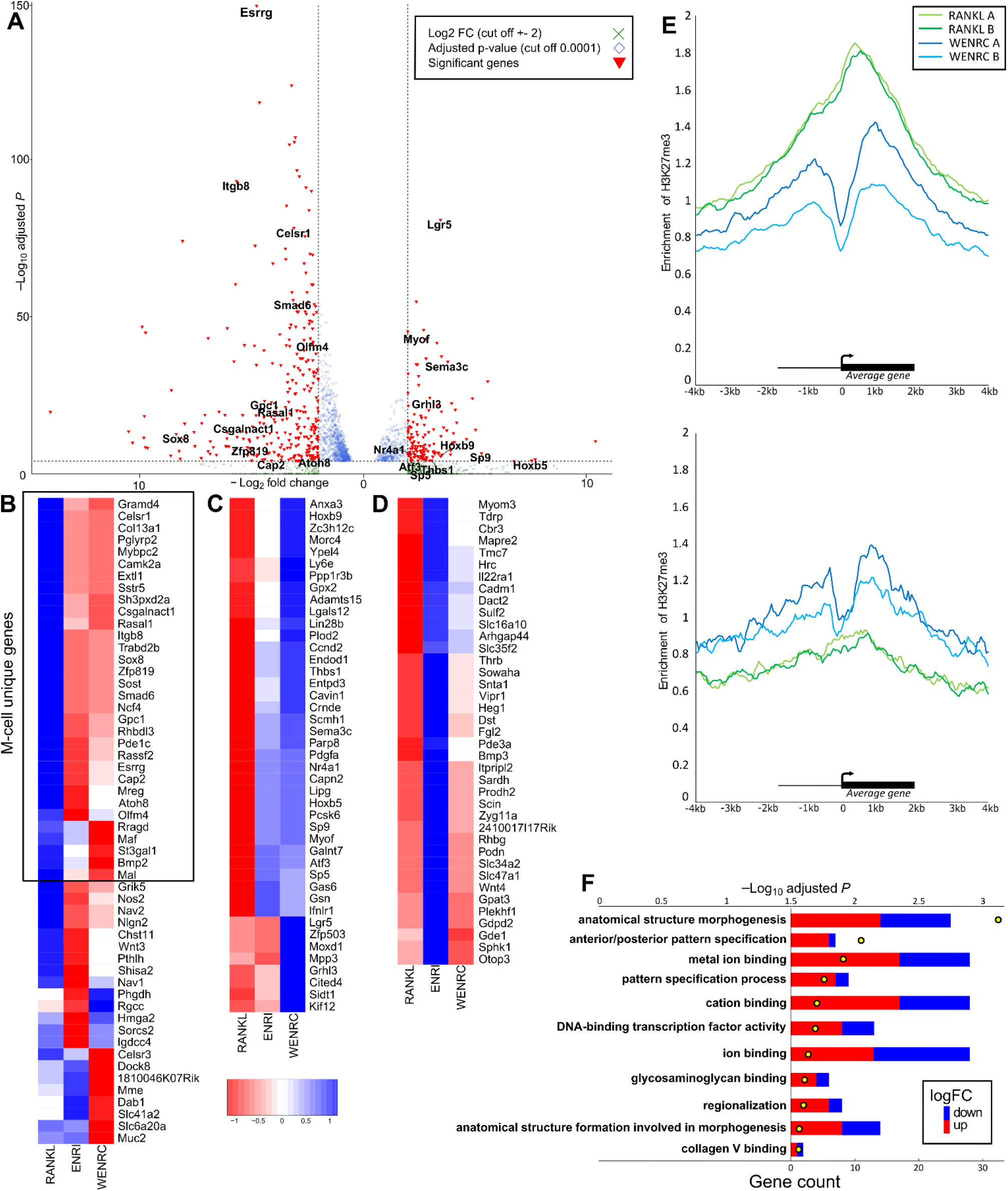
PRC2 regulated genes during the M cell differentiation. A) Differentially expressed genes during M cell differentiation detected with Gro-Seq. Signal is depicted by volcano plot comparing organoids before and after RankL treatment. X-axis and Y-axis indicate the log2 fold change and –log10 adjusted P-value. Differentially expressed genes are marked (GRO-Seq with log2 fold change cutoff at +-2 and P-value < 0.0001). Upregulated genes from ENRI conditions were removed to show only RankL specific regulation. B) Genes upregulated in M cells compared to stem cells and enterocytes. Heatmap of differentially expressed (GRO-Seq with log2 fc cutoff at +-2 and P-value < 0.0001) and H3K27me3 regulated (log2 fc +-2, P-value < 10-6) genes in RankL, ENRI and WENRC treated organoids showing centered log2 fold change. C) Genes downregulated by H3K27me3 in M cells and enterocytes compared to stem cells. D) Genes downregulated by H3K27me3 in stem cells and M cells compared to enterocytes. E) Composite enrichment analysis of H3K27me3signal density +-4000 bases around transcription start sites in genes specifically silenced by PRC2 in M cells (above) and specifically expressed in M cells (below). F) Gene ontologies enriched (P < 0.05) in differentially expressed PRC2-target genes between M cells and stem cells (GRO-Seq log2 fc +-2, P-value <0.0001 and H3K27me3 log 2 fc +-2, P-value < 10-6) in molecular function and biological process.

### Esrrg is expressed in M cells and induced by Rank-RankL signalling

Of the six transcription factors that were specifically expressed in M cells in PRC2-dependent manner (Fig. 2 A&B), Estrogen related receptor gamma (Esrrg) turned up as one of the most highly expressed PRC2-regulated gene during M cell differentiation. (log2 fold changes –6.64 RankL vs. ENRI and – 4.76 in RankL vs. WENRC comparisons) (Fig. 3A). Immunohistochemistry analysis for Esrrg in Peyer’s patch shows that Esrrg was localized in the FAE cells (Fig. 3B). RNA was isolated from FAE isolated from Peyer's patch and villus epithelium and the RT-qPCR analysis confirmed that the Esrrg was significantly enriched in FAE (GP2 as a marker) when compared to villus epithelium (Fig 3C). To ascertain that Esrrg is a novel RankL-induced M cell gene, RT-qPCR analyses was performed for organoids before and after 4 days of RankL treatment. The expression of Esrrg, together with M cell marker Gp2, was significantly induced with RankL treatment (Fig. 3D). To confirm that Esrrg expression was specifically regulated by Rank-Rankl signalling, we generated Rank-deficient mouse intestinal organoids using Lenti-V2 CRISPR/Cas9 system and found that in Rank-deficient organoids, RankL treatment did not induce the expression of Esrrg (Fig. 3E). The Rank KO was validated by immunoblot analysis (supplemental Fig 2A)

**Figure 3.**
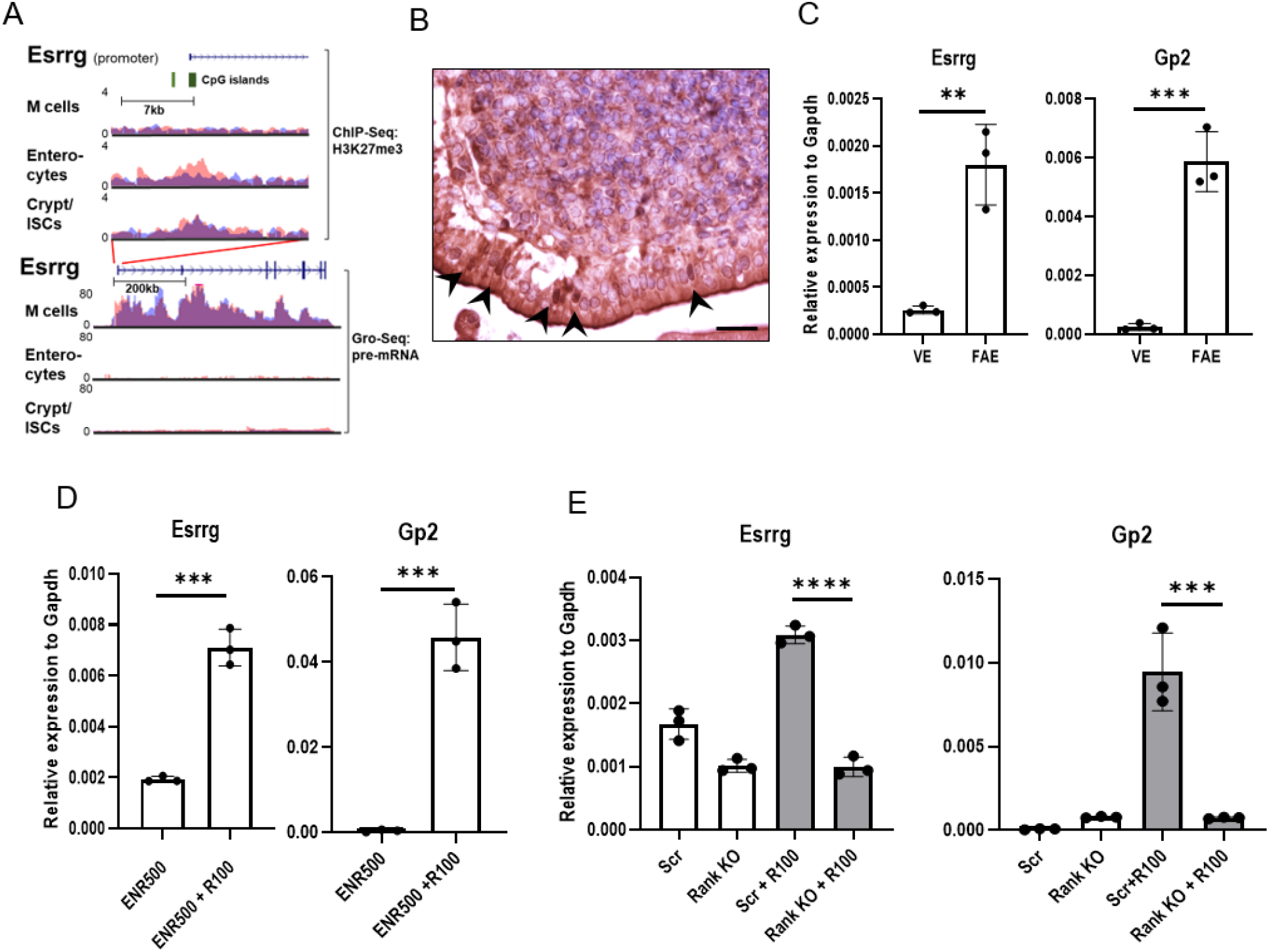
Esrrg is expressed in FAE in Peyer’s patches and is dependent on Rank-RankL signalling. A) H3K27me3 occupancy at CpG islands spanning the promoter and first exon of the Esrrg gene in organoids treated with RankL (M cells) or inhibited with IWP2 (Enterocytes) or treated with Wnt3a and Chir99021 (Crypt/ISCs). Below, pre-mRNA expression of Esrrg in organoids treated as above (y-axis: normalized tag count, ENR500 = R-spondin 500ng/ml, R100 = Rankl 100ng/ml). B) Section of PP from wild-type mice stained with Esrrg antibody. Arrowheads indicating Essrg expression in the nuclei of M cells in FAE. C) RT-qPCR analysis of Esrrg and Gp2 in the FAE and VE from C57BL/6JRj mice (*n* = 3 from wild-type mice). D) Organoids generated from wild-type mice were stimulated with 100ng of RankL for 4 d. Esrrg and Gp2 expression was examined by quantitative PCR analysis. E) Rank KO organoids and Scrambled organoids generated by lentiCRISPR v2 were incubated with RankL for 4 days, Esrrg and GP2 expression was analyzed by quantitative RT-qPCR. In (C-E) unpaired two-tailed Student's *t* test was performed for three independent experiments, ****, P < 0.0005; ***, P < 0.005; **, P < 0.01.

### Non-canonical NF-κB activation is necessary and sufficient for Esrrg expression

Rank-RankL signaling has been previously shown to activate canonical as well as non-canonical NF-κB pathways. LTβR signaling was implicated in inducing both classical p50–RelA and non-canonical p52–RelB heterodimers in Peyer’s patch (Yilmaz et al.,2003). Mouse intestinal organoids were grown for 3 days in the presence of LTα1β2, the ligand of LTβR, and we observed that, similar to Spi-B, Esrrg expression was also significantly increased (Fig 4A). To identify if canonical NF-κB specifically had a role in Esrrg expression, mouse organoids were grown in the presence and absence of RankL and SC-514, a specific inhibitor of IκB kinase-β (Kishore et al., 2003). We found that inhibiting canonical NF-κB with SC-514 completely abrogated the expression of Esrrg (Fig 4B), similarly as was reported for the Spi-B (Kanaya et al., 2018). Both, p50-RelA and p52-RelB overexpression led to increased expression of Esrrg (Fig 4C). It was previously demonstrated that p50/RelA (canonical NF-κB) directly targets the transcription of RelB (Bren et al., 2001). The treatment of p52/RelB overexpression organoids with SC-514 could not suppress the activation of Esrrg (Fig. 4D). To conclude, these data indicate that non-canonical NF-κB is necessary and sufficient to induce Esrrg.

**Figure 4.**
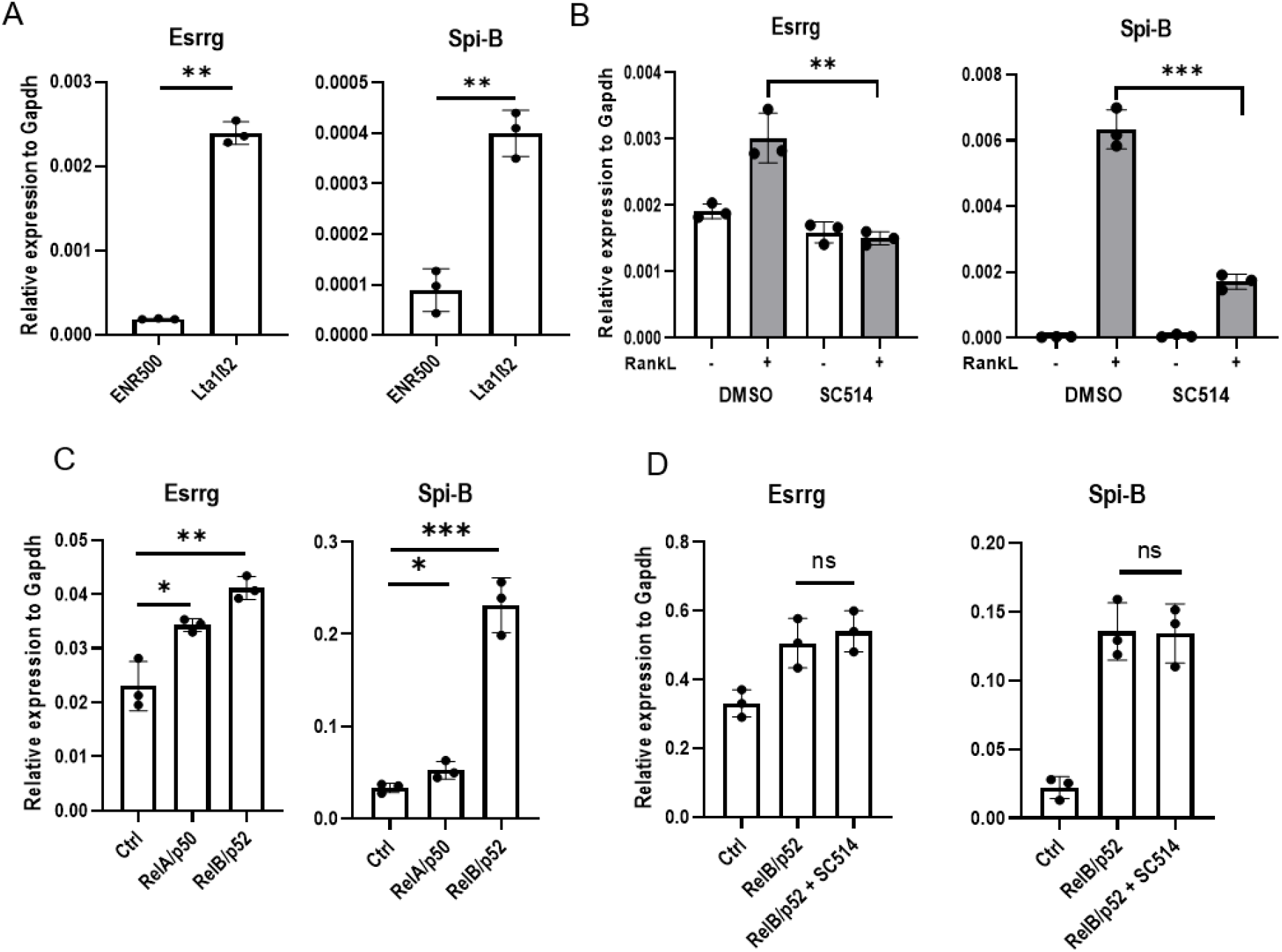
Esrrg expression is induced by RelB/p52 activation. A) Ltα1β2 prominently upregulated the expression of Esrrg in organoids. Organoids from C57BL/6JRj mice were stimulated with LTα1β2 for 3 d, and the gene expression was analyzed by quantitative PCR. B) Organoids from wild-type mice were stimulated with RankL for 3 d in the absence or presence of 125μM SC-514. Gene expression was analyzed by quantitative PCR. C) Organoids were transduced to express classical and non-canonical NF-kβ and the expression of Esrrg and Spi-B (control) are represented in relative to Gapdh D) Organoids expressing p52 and RelB in the presence of SC-514 for 3 d, and the expression of Esrrg and Spi-B was analyzed by quantitative PCR. Values in all are presented as the mean ± SD and unpaired two-tailed Student’s *t* test was performed, n=3; n.s, not significant; **, P < 0.01; ***, P < 0.005.

### Esrrg expression is required for Sox8 activation and maturation of M Cells

Given that Esrrg was prominently expressed in Peyer’s patches and regulated by non-canonical Nf-kb expression, we sought to see if abolition of Esrrg had any effect on M cell differentiation and development. To investigate this, mouse intestinal organoids deficient in Esrrg were generated by LentiCRISPR V2 CRISPR-Cas9 genome editing. Targeting of Esrrg by the gRNA selected (Fig. 5A) resulted in a significant reduction in expression of Esrrg protein (Fig 5B). Esrrg deficient organoids were grown in the presence and absence of RankL for 3 days. RT-qPCR analysis revealed that Gp2 was nearly absent and Sox8 expression was significantly reduced in Esrrg targeted organoids (Fig. 5C). Gp2 and Sox8 immunostaining in Esrrg targeted organoids showed an absent and reduced expression respectively (Fig 5D). However, expression of Spi-B and Tnfaip2 remained unaffected in the Esrrg deficient organoids (Fig 5C and 5E). Early M cell differentiation markers such as CCL20, CCL9 and MarksL1 were significantly affected by the lack of Esrrg as well (Fig 5E). Aif1, which is a regulatory gene for transcytosis in M cells was also found to be severely affected by lack of Esrrg expression (Fig 5E). Our observations show that Esrrg is required for the expression of Sox8 and for the early markers as well as late maturation steps in M cell differentiation. To validate for off-target, we knocked out Esrrg with a second guide RNA, comparable results were observed (Supplementary Figure 1A and 1B).

**Figure 5.**
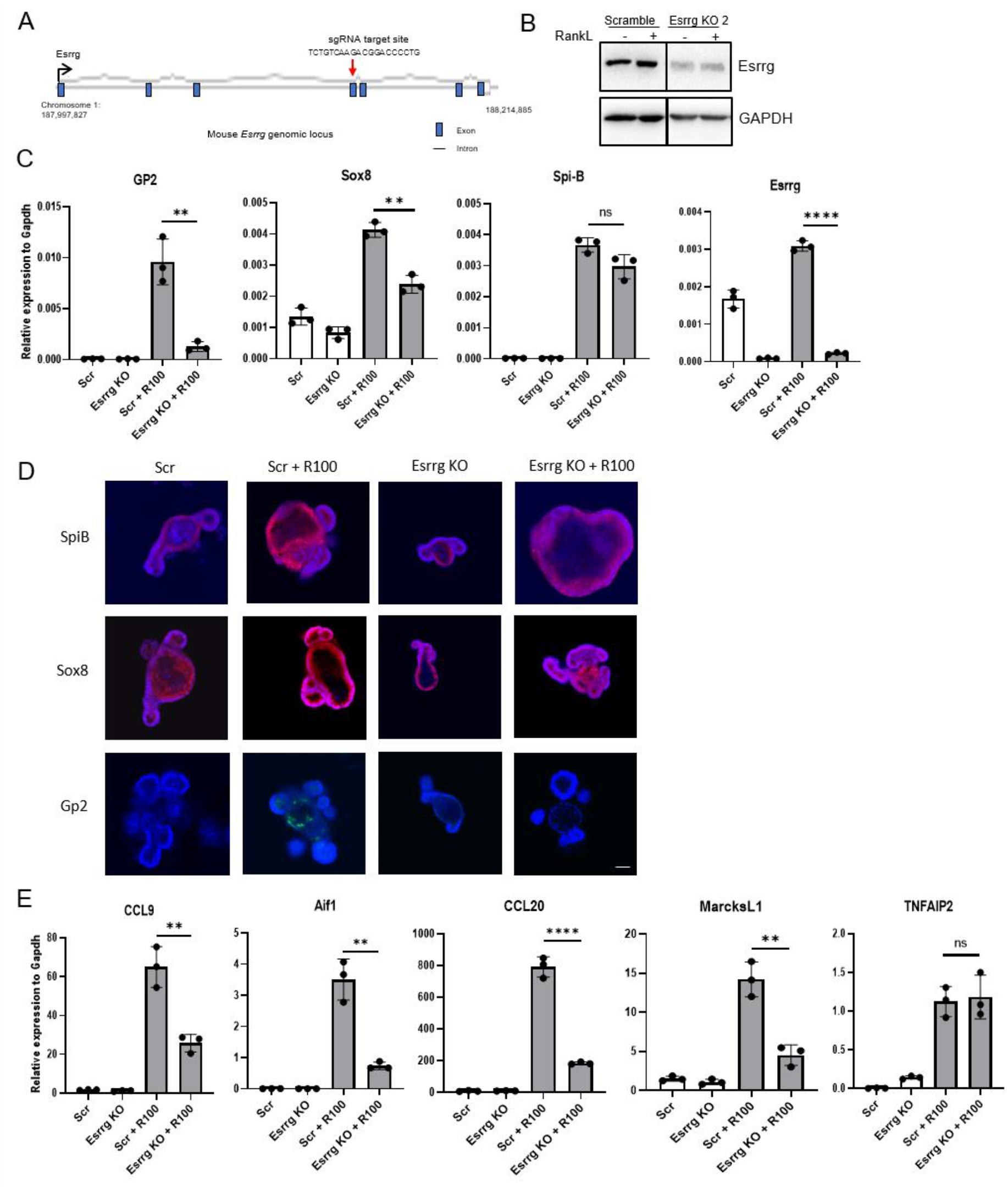
Abolition of Esrrg impairs Sox8 activation and the functional maturation of M cells. A) Schematic representation of Esrrg knockout design by CRISPR-Cas9 genome editing in mouse intestinal organoids. Exon, intron, and genomic position are indicated. B) Esrrg protein expression in Esrrg KO and Scrambled cells generated by lentiCRISPR v2 genome editing in C57BL/6JRj intestinal organoids. Organoid lysates were analyzed by Western blot. C) qPCR analysis of M cell associated genes expressed in in vitro culture stimulated by RankL for 4 days. D) Immunostaining images for Spi-B (red), Sox8 (red) and Gp2 (green) in Scrambled organoids with and without RankL 100ng for 4 days and Esrrg KO with and without RankL 100ng, Bars, 100 μm. E) qPCR analysis of early markers of M cell associated genes expressed in vitro in the presence of RankL for 4 days. Values in all are presented as the mean ± SD. Unpaired two-tailed Student’s t test, n=3, ns, not significant; **, P < 0.01; ****, P < 0.0005.

### Esrrg acts upstream of Sox8 expression

Spi-B and Sox8 were found to be two key transcription factors involved and essential for M cell differentiation and regulation of expression of other M cell associated genes (de Lau et al., 2012; Kanaya et al., 2012; Kimura et al., 2019; Sato et al., 2013). Spi-B and Sox8 mutant mice lacked Gp2+ mature M cells and were unable to transcytose antigens. However, it was observed that Spi-B was dispensable to the expression of Sox8 even though Spi-B expression was moderately reduced in the Sox8 mutant mice (Kimura et al., 2019). To investigate if Esrrg is affected by knockout of Sox8, RNA was isolated from FAE of Peyer’s patch from Sox8 +/+ and Sox8 -/- mice. RT-qPCR analysis of FAE’s from PP’s of Sox8 +/+ and Sox8 -/- demonstrated that the expression of Esrrg remained the same and even higher in Sox8 -/- mice (Fig. 6A). Organoids isolated from Sox8 +/+ and Sox8 -/- mice were treated with RankL and expression of Esrrg expression was analysed by RT-qPCR and Western blot. The analysis showed Esrrg expression was intact and similar to the in vivo data (Fig. 6B and 6C). This suggests that Esrrg acts upstream of Sox8 and could play a role in the activation of Sox8. Next, we explored how Esrrg was affected by Spi-B. Spi-B deficient organoids were generated by LentiCRISPR V2 genome editing and grown in the presence and absence of RankL. qPCR analysis showed that Esrrg was moderately affected by the lack of Spi-B (Fig 6D). Western blot analysis also indicated that Essrg was expressed less in Spi-B deficient organoids (Fig 6E) Spi-B Ko organoids were validated by immunoblot analysis (supplemental Fig 2B).

**Figure 6.**
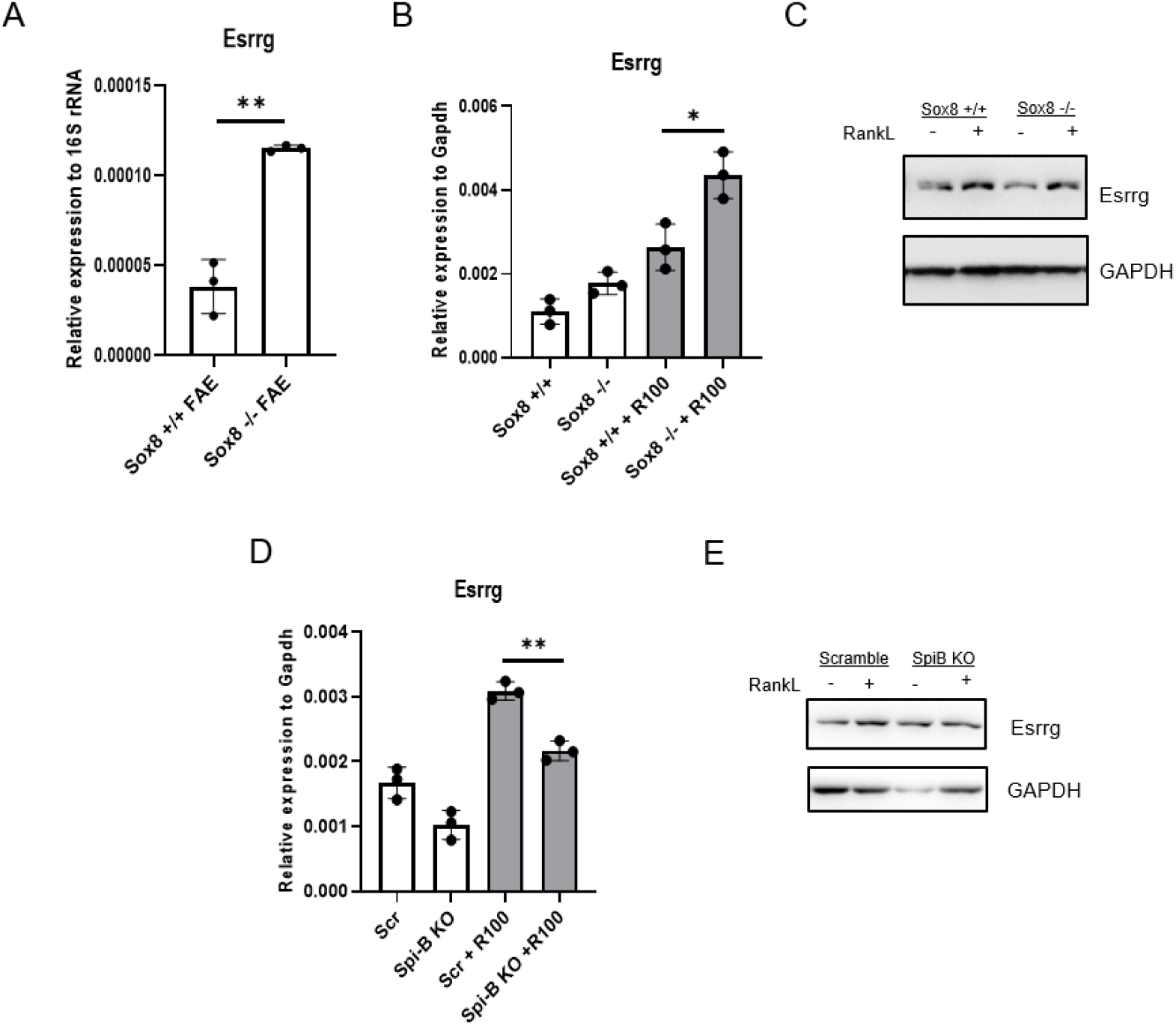
Esrrg expression and its relation to other M Cell developmental markers Spi-B and Sox8. A) Esrrg expression was unaffected by lack of Sox8 expression. qPCR analysis of Esrrg and Gp2 in the FAE and VE from Sox8+/+ and Sox8-/- mice. B) Organoids generated from Sox8 +/+ and Sox8 -/- mice were stimulated with and without RankL for 4 d. Esrrg expression was examined by quantitative PCR analysis. C) Organoids isolated from Sox8 wild-type and Sox8 KO mice were lysed and analyzed by Western blot for Esrrg expression. D) qPCR analysis of Esrrg in a Spi-B knockout intestinal organoids by lentiCRISPR v2. E) Organoid lysates for Esrrg in Scrambled and SpiB KO organoids were analyzed by Western blot. Values in all are presented as the mean ± SD. Unpaired two-tailed Student’s t test, n=3, ns, not significant; *, P<0.05; **, P < 0.01.

### Overexpression of Esrrg is not sufficient for GP2+ M cells but Esrrg agonist augmented GP2 expression

Spi-B expression in Sox8 KO mice and Sox8 expression in Spi-B KO mice did not lead to a mature GP2 M cell phenotype in either of these mice. As we found that the expression of Gp2 was dependent on Esrrg (Fig. 5C) we sought to investigate if the overexpression of Esrrg alone could lead to upregulation of Sox8 or Gp2 expression. Esrrg cloned into CSII-CMV-MCS-IRES2-Bsd overexpression vector was transduced into mouse intestinal organoids. RT-qPCR analysis showed that Esrrg alone was not adequate enough to induce Gp2 or other M cell specific transcription factor such as Spi-B or Sox8. (Fig 7A). Esrrg is an orphan nuclear receptor without known natural ligands. However, 4-hydroxytamoxifen (4OHT) has been shown to bind and inhibit Esrrg activity and phenolic acyl hydrazones-GSk4716 has been identified as an agonist that enables the activation of other co-activators and other downstream targets (Coward et al., 2001.; Zuercher et al., 2005). The antagonist 4OHT along with RankL induced a similar result to the lack of Esrrg protein, Spi-B remained unaffected, whereas Sox8 and Gp2 were significantly affected (Fig 7B). Organoids treated with agonist GSK4716 and RankL did not show significant increase in Sox8 or Spi-B expression however, Gp2 expression was slightly augmented with GSK4716 (Fig 7C).

**Figure 7.**
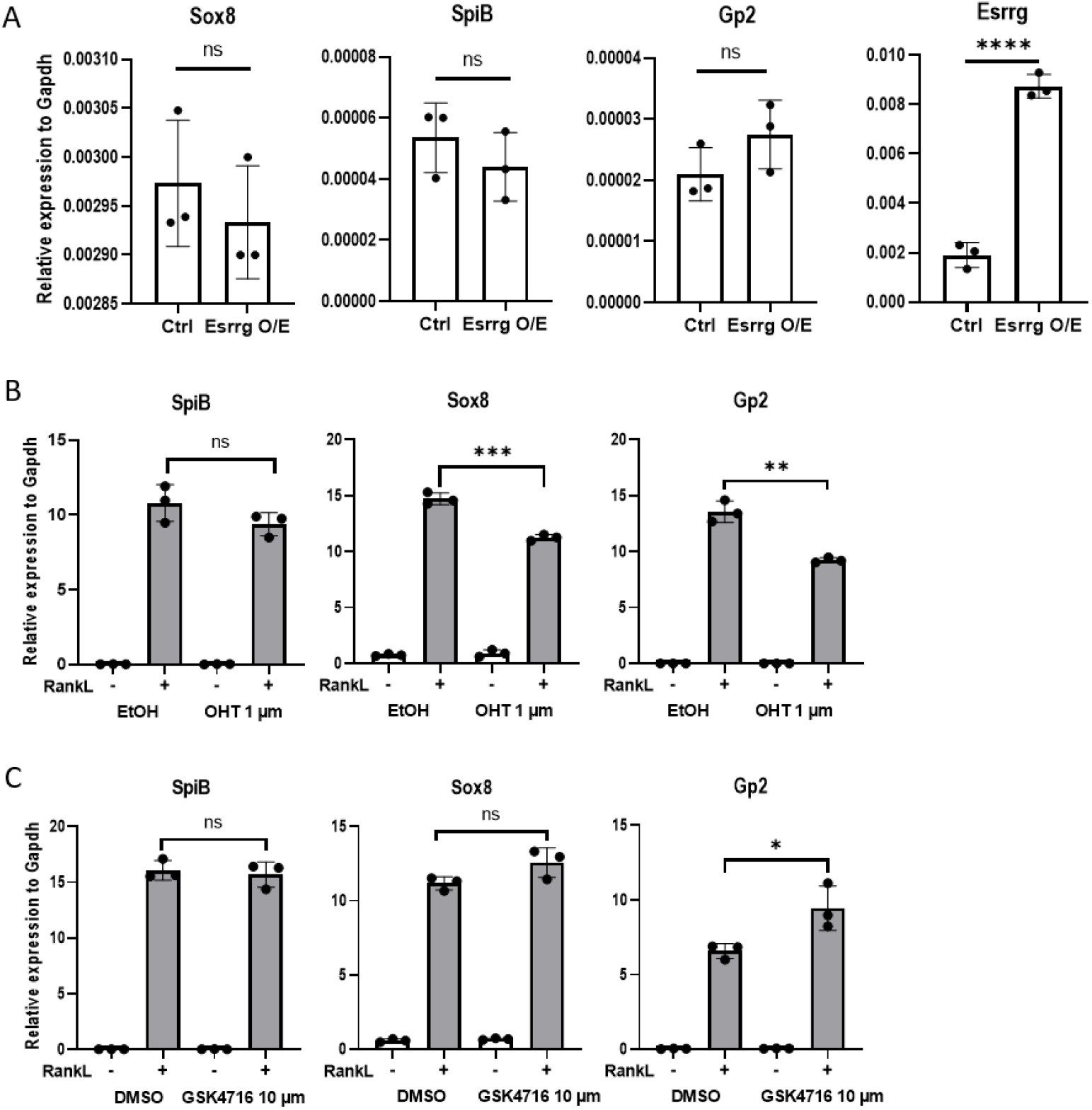
Esrrg alone is not sufficient for maturation of GP2+ M cells. A) Intestinal organoids were dissociated and transduced by lentivirus encoding Esrrg. qPCR analysis of Spi-B, Sox8 and GP2 showed no significant changes. B) Organoids were grown in the presence and absence of 100ng/ml Rankl and 1um of tamoxifen-antagonist of Esrrg. Spi-B, Gp2 and Sox8 expression was analyzed with RT-qPCR. C) Intestinal organoids from mouse were grown in the presence and absence of Rankl 100ng/ml and 10um GSK 4718-agonist of Esrrg for 3 days. Spi-B, Gp2 and Sox8 analyzed with RT-qPCR. Values are presented as the mean ± SD. Unpaired two-tailed Student’s t test, n =3. n.s., not significant. *, P < 0.05; **, P < 0.01; ***, P < 0.005.

## Discussion

Our data reveals the PRC2-regulated genes during the differentiation of intestinal microfold cells. As PRC2 is the master regulator of development it is very likely that many of the identified PRC2 targets, that are specifically induced in M cells, contributes to the maturation of this cell type. Transcription factors are usually expedient to development and we identified six PRC2 regulated transcription factors specifically upregulated after the RankL-induced M cell differentiation. Amongst those was previously identified transcription factor Sox8 (Kimura et al, 2019) which we also here showed to be necessary to M cell differentiation. As Esrrg was clearly the most highly induced PRC2-target gene during the M cell differentiation, we pursued to study it in more detail in intestinal organoids and showed that it is indispensable to the maturation of intestinal stem cells into Gp2+ M cells. Esrrg is a member of the ESRR nuclear receptor family which also includes ESRRA and ESRRB (Giguère, 2002). This subfamily of orphan nuclear receptors has shown to share target genes, coregulatory ligands, and sites of action with ERs. Esrrg was implicated to control macrophage function indirectly through regulation of intracellular iron. In response to *Salmonella typhimurium* infection, hepatic expression of the hormone hepcidin is upregulated by ERRγ downstream on IL-6 signalling (Bren et al., 2001; D. K. Kim et al., 2014). However, no prior data about the expression and function of Esrrg in M cell and M cell induced transcytosis of antigens exist. We defined the specific expression of Esrrg by M cells in mouse FAE’s and how its expression was upregulated by induction of RankL and was under the influence of RankL-Rank pathway. The loss of Esrrg led to lack of expression of GP2 receptor in Esrrg KO organoids which is characteristic of a mature M cell and lack of Gp2 in M cell has been shown to result in attenuation of antigen sampling and transcytosis. This distribution of Esrrg along with the phenotype of Esrrg KO organoid highlights a critical role for Essrg in the maturation of functional M cells

RelB/p52 was shown to upregulate Spi-B (Kanaya et al., 2018). Similarly, here we show that Esrrg is also regulated by the activation of the non-canonical NF-κB pathway. We believe that the expression of Esrrg and Spi-B is regulated downstream parallelly by RankL-Rank-RelB/p52 signalling (Fig. 8). Esrrg has been shown to behave as a constitutive activator of transcription. (Matsushima et al., 2007). Here we demonstrate that Esrrg is needed for the activation of Sox8. Kimura et al previously showed how Sox8 was indispensable for the expression of Gp2 and Sox8 KO mice showed a decrease in uptake of antigens and a significant decrease of IgA+ immunoglobulins. Sox8 was also shown to bind directly to Gp2 promoter along with SpiB. The significant decrease in Sox8 expression could explain as to why Esrrg KO organoids were not able to activate Gp2. Sox8 expression alone cannot lead to an increase in Gp2 expression because enhancer activation by SOX proteins require DNA binding partners specific for each member of the SOX family. These DNA binding partners aid in stabilizing the SOX family to their target regions (Kamachi et al., 1999). This suggests that Esrrg activated Sox8 requires another molecule downstream of Spi-B or through another pathway to bind to Gp2 to induce its expression, however further exploration is required to confirm this. Esrrg overexpression alone did not lead to expression of Sox8 confirming the need of a ligand and/or other factors to activate downstream targets.

**Figure 8.**
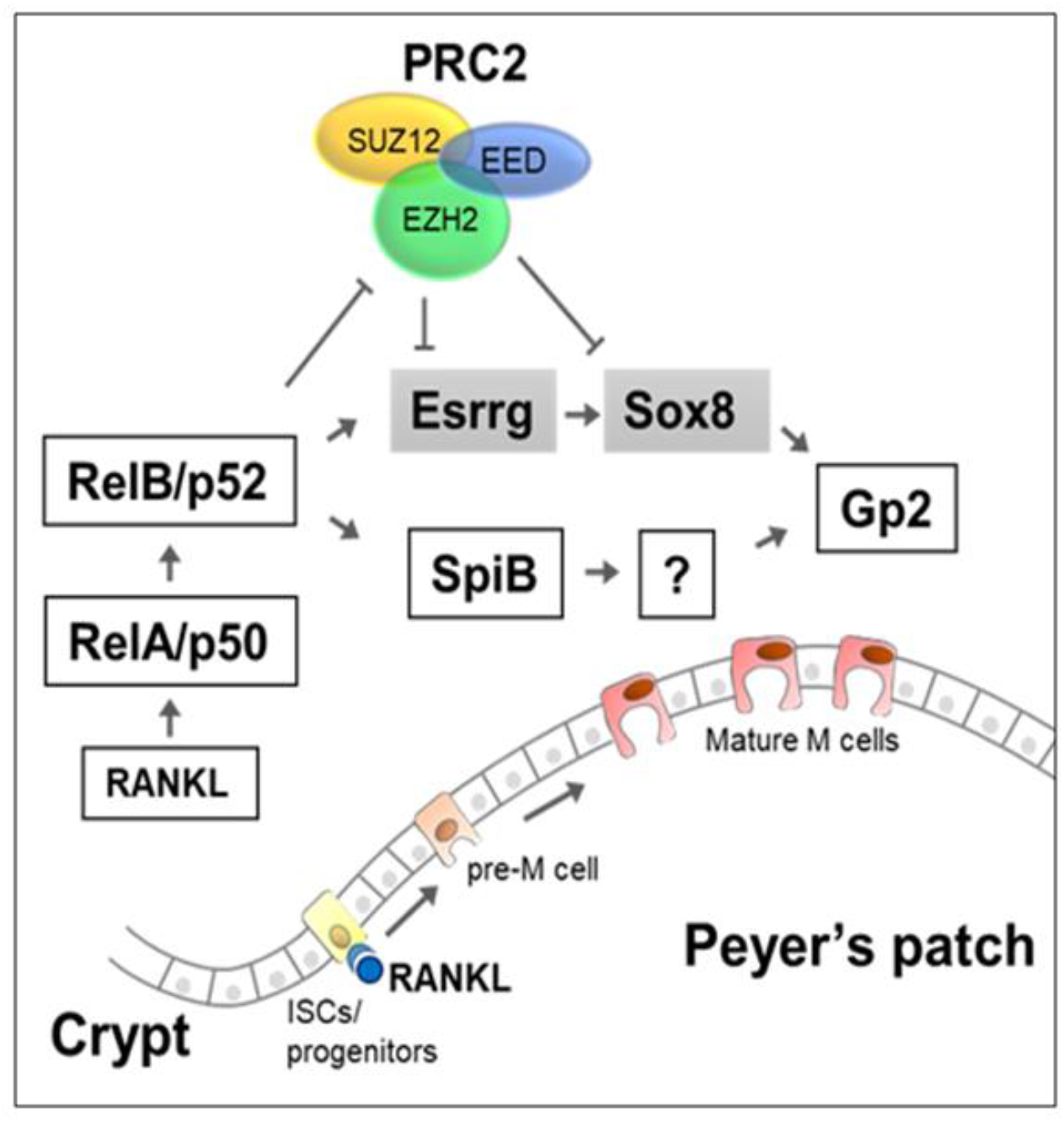
PRC2 regulates the differentiation of M-cell. In the absence of Rank-RankL signaling, Esrrg and Sox8 genes are repressed by PRC2 in the intestinal epithelium. When migrating progenitors with Rank receptors binds to RankL in Peyer’s patches, this induces NF-κB signaling leading to loss of H3K27me3 from the gene promoters and activation of Essrg and Sox8. Sustained expression of Sox8 and GP2 and differentiation of M cells is dependent on Esrrg.

A major portion of our investigation of Esrrg in M cells was conducted in vitro and Esrrg ablation in mouse models is required to further ascertain its role. Interestingly, Sox8 deficiency did not affect early M cell markers, however, Esrrg KO organoids showed a drastic decrease in expression of early M cells markers such as CCL9 and MarcksL1. Mature marker Gp2 receptor expression was significantly decreased as well. The loss of Aif1 expression in Esrrg knockout means that transcytotic capacity of the M cells would be affected as well. In the Sox8 KO mice, we observed that Esrrg expression was still intact and even observed to have a higher expression in vivo and in vitro suggesting that Esrrg was not affected by the absence of Sox8 and acts upstream of it. The significance of Esrrg was further confirmed with our antagonist and agonist experimental study. Even though tamoxifen is known to trigger multiple signalling pathways in the cell, it has been identified as an antagonist for Esrrg receptor, tamoxifen 1um was enough to significantly decreased the expression of Gp2 and Sox8 similar to the Esrrg KO organoids. The treatment of intestinal organoids with RankL and Esrrg agonist GSK4716 augmented the expression of Gp2 when compared to organoids with just RankL treatment. This could be because GSK4716 binds to Esrrg and activates several other unknown downstream targets that combine to attenuate Gp2 expression. Esrr family members are known to be orphan receptors meaning they might not need a ligand for its function, or the ligand remains unknown. Overexpression of Esrrg did not lead to an increase in Sox8 expression or other transcription factors necessary for mature M cell differentiation presumably because a specific ligand might be necessary. However further studies are needed to prove this.

In conclusion, we identified several previously unknown PRC2 regulated genes implicated in M cell differentiation. One of the genes, we identified, Esrrg, is as a key transcription factor required for the functional development and M cell differentiation which is pertinent for constant surveillance of the mucosal lining of the gastrointestinal tract. We believe that the further exploration of other activators of Gp2 will lead to better elucidation of M cell maturation and antigen transcytosis. This will create potential to provide strategic innovation in support of mucosal/oral vaccine advancement.

## Materials and Methods

### Animals

C57BL/6JRj mice were purchased from Janvier labs and were maintained with constant breeding. The *Bac-Cre-ERT2;Sox9f/f;Sox8-/-;Rosa26Eyfp* mice were a gift from Raphael Jimenez (Granada, Spain). These mice were backcrossed with C57BL/6JRj to isolate Sox8-/- allele. Sox8+/- were bred to obtain Sox8-/- and littermate controls: Sox8+/− and Sox8 +/+. F1-4 mice were used for gene or protein expressions. Genotyping of the wild-type, heterozygous, and deleted alleles was carried out by PCR with the following primers: F1, 5′-GTCCTGCGTGGCAACCTTGG-3′; R1, 5′- GCCCACACCATGAAGGCATTC-3′; F3, 5′-TAAAAATGCGCTCAGGTCAA-3′. Conventional conditions were observed for the maintenance of these mice at the pathogen free animal facility of the faculty of Medicine and Health Technology. All animal experiments were approved by the Finnish National Animal Experiment Board (Permit: ESAVI/5824/2018).

### Intestinal Organoid culture

Intestinal crypt isolation and culture techniques were observed as previously described by Sato et al 2011 and de Lau et al 2012. Mouse duodenum were cut longitudinally, and the villi were gently scraped off with 2 glass slides. After a couple of PBS washes, they were cut into 2mm pieces and pipetted up and down 5 times in 15ml PBS with a 10ml pipette, this step was repeated 3 times with fresh PBS. The pieces were incubated in 10mM EDTA in PBS for 20 minutes rocking at room temperature. The pieces were vigorously suspended in cold PBS and the mixture was strained through a 70-μm cell strainer (Fisher Scientific). This mixture was enriched to crypt fraction through centrifugation at 150 × g for 5 minutes. The enriched crypts were embedded in Matrigel (Corning) and 30ul were plated on a 24 well plate. Crypts were cultured in an optimal medium consisting of advanced DMEM/F12 (Thermo Fisher Scientific) that contained HEPES (10mM, Sigma-Aldrich), Glutamax (2mM, Thermo Fisher Scientific), Penicillin-streptomycin (100U/ml, Sigma-Aldrich), B-27 supplement minus Vitamin A (Thermo Fisher Scientific), N-2 supplement (Thermo Fisher Scientific), *N*-acetylcysteine (1 mM; Sigma-Aldrich), recombinant murine EGF (50 ng/mL; Thermo Fisher Scientific), recombinant murine Noggin (100 ng/mL; PeproTech), recombinant mouse R-spondin1 (1 μg/mL; R&D Systems). Media were changed every 2 days. For M cell differentiation, recombinant mouse RankL (100ng/ml, Peprotech) was added to the media and incubated for 4 days. PRC2 was inhibited by addition of 5μm of Ezh2 inhibitor (EI1) (CAS 1418308-27-6 Calbiochem Chemicals). For activation of NF-κB, human LTα1β2 (1 μg/ml, R&D) were added into organoid cultures. Restriction of IκB kinase-β activity was achieved by adding, SC-514 (125 μM Calbiochem) for 3 days (Akiyama et al., 2008). (Z)-4-Hydroxytamoxifen (1μm, Sigma-Adrich) was used as an antagonist of Esrrg and GSK 4716 (10 μm, Sigma-Aldrich) was used as an agonist of Esrrg.

### ChIP-Seq analysis

Intestinal organoids (in ENR500, ENRI and RankL culturing conditions) were isolated from Matrigel with Cell Recovery Solution (Corning). This was followed by washes with cold PBS and dissociation into single cell suspension using TrypLE Express (Thermo Fisher Scientific) and counted. 10 × 10^6 of cells of each condition were crosslinked with formaldehyde, after which nuclei were isolated with Lysis buffer 1, 2 and 3 as described (Lee et al., 2006) and sonicated with a Covaris S220 ultrasonicator (Woburn, MA). The resulting nuclear extract was incubated with Dynal protein G beads which were preincubated with 5μg of H3K27me3 or H3 antibody (ab6002 and ab1791, Abcam) respectively at 4°C overnight. After washing and elution of bound complexes from the beads, crosslinks were reversed by heating to 65°C. IP and input DNA were then purified by a treatment with RNAse A, proteinase K, and phenol:chloroform extraction. NEBnext UltraDNA-library preparation kit for Illumina (NEB, Ipswich, MA) was used to construct libraries from IP and input DNA and subjected to 50 bp single-end read sequencing with Illumina Hiseq 2000 at EMBL Genecore, Heidelberg, Germany.

### Gro-Seq analysis

ENR500 and RankL treated organoids were harvested (as in ChIP-Seq) and GRO-Seq was performed for equal number of isolated nuclei. Nuclei extraction and run on reaction was performed as described (Core LJ et al., 2008). For each replicate, 3 million cells were suspended to a final volume of 80-200 μl of freezing buffer. Trizol LS (Life Technologies) was used to extract RNA and fragmented for 13 mins in 70°C using RNA Fragmentation Reagents (Life Technologies) and later purified by running through RNase-free P-30 column (Bio-Rad, Hercules, CA). PNK was used to dephosphorylate RNA for 2 hours (New England Biolabs, Ipswich, MA) followed by heat-inactivation. 65 μl of blocking solution was added (5x volume of 0.25xSSPE, 1 mM EDTA, 37.5 mM NaCl, 0.05% Tween-20, 0.1%PVP and 0.1% ultrapure BSA for 1 hour in RT), the anti-BrdU bead slurry (Santa Cruz Biotech) suspended in 500μl of binding buffer (0.25xsSSPE, 1 mM EDTA, 37.5 mM NaCl 0.05% Tween-20) was used to purify the dephosphorylated reaction. After binding for an hour in RT, the beads were washed 2x with binding buffer, 2x with low salt buffer (0.2xSSPE, 1mM EDTA, 0.05% Tween-20), 1x with high salt buffer (0.2xSSPE, 1mM EDTA, 135 mM NaCl0.05% Tween-20) and lastly 2x with TE-buffer (1xTE, 0.05% Tween-20). The elution of the RNA was completed with 130μl of elution buffer (50 mM Tris-HCl pH 7.5, 150 mM NaCl, 0.1% SDS, 1mM EDTA, and 20 mM DTT) followed by ethanol precipitation overnight. All buffers were supplemented with SUPERase ln (2μl /10 ml; Life Technologies). Library preparations were performed the next day as previously reported by Kaikkonen Mu et al 2014. The library was amplified with 14 cycles and the final product of 190-135 bp was extracted from a 10% TBE gel. The DNA was purified from the gel using the Gel extraction Kit (Thermo) and eluted in TE buffer (TE 0.1% Tween + 150 mM NaCl). ChIP DNA clean & Concentrator Kit (Zymo Research Corporation, Irvine, CA) was used to purify the library, the DNA was quantified with the Qubit fluorometer and sequenced with Illumina HiSeq 2000 at EMBL Genecore, Heidelberg, Germany.

### ChIP- and Gro-Seq data analyses

Analyses were performed as described by Core LJ et al 2008, Kaikkonen Mu et al 2014, and Oittinen et al 2016. The data has been deposited in the NCBI Gene Expression Omnibus database (GSE157629). Gro- and ChIP-seq data for the individual genes are shown in Supplementary tables 1 and 2 respectively.

### Immunohistochemistry and immunofluorescence

Peyer’s patches from the ileum were isolated and washed with cold PBS and embedded into paraffin blocks. Sections from the blocks were rehydrated and washed with PBS. After incubation with 1% PBS/BSA supplemented with 5% normal donkey serum for blocking. Antigen retrieval was processed with citrate buffer, pH 6.0 (121°C for 5 min) and stained overnight at 4°C for Esrrg (abcam, ab49129), Gp2 (MBL, D278-3) antibodies. This was followed by anti-Rabbit secondary for Esrrg and Anti-Rat secondary for Gp2. The sections were examined with a light microscope. For whole-mount immunostaining, crypt organoids were plated in an 8-well chamber and cultured for 4 days after which they were fixed with 4% PFA, followed by permeabilization with 0.1% Triton X-100. The organoids were stained with the following primary antibodies overnight at 4°C: rabbit anti-Spi-B (Spi-B (D3C5E), CST, 14223), rabbit anti-Sox8 (abcam, ab221053), rat anti-Gp2 (MBL, D278-3). This was followed by incubation with secondary antibody anti-rabbit Alexa fluor 568 for Spi-B and Sox8 and anti-Rat Alexa fluor 488 for Gp2. Cells were analyzed with Nikon A1R+ Laser Scanning Confocal Microscope after mounting with ProLong Diamond with Dapi mounting solution (Molecular Probes P36962).

### Isolation of villous epithelium and follicle associated epithelial cells

Villous Epithelium (VE) and Follicle Associated Epithelium (FAE) were prepared by isolating illeal PP’s and small pieces of ileum from the intestine. These pieces were washed in cold PBS and later incubated in 30mM EDTA, 5mM DTT in PBS and gently shaken in ice on a rocker for 20 minutes. After which surrounding epithelial cells were peeled off from lamina propria and PP’s. FAE was carefully cleaned off from surrounding VE tissues with a 26-gauge needle under a stereo microscope.

### CRISPR–Cas9 gene editing of intestinal organoids

Guide RNA’s for Rank, Spi-B and Esrrg were designed with CRISPR design tool (http://crispr.mit.edu). (Shalem, O. et al. *Science* 343, 84–87 (2014)). The guides were cloned into lentiCRISPR v2 vector (Addgene, 52961). The cloned vector was transfected into 293FT cells (ThermoFisher R7007) and the supernatant was collected at 48 hours and concentrated with Lenti-X concentrator (Clontech). The 293FT cell line was found to be negative for mycoplasma. Cultured intestinal organoids were grown in ENCY (EGF, Noggin, Chir-99021 and Y-27632) 2 days prior to transduction. Organoids were dissociated into single cells mechanically along with TrypLE Express (Thermo Fisher Scientific) supplemented with 1,000 U/ml DnaseI for 5 min at 32 °C. The single cell suspension was washed once with Advanced DMEM and resuspended in transduction medium (ENR media supplemented with 1mM nicotinamide, Y-27632, Chir99021, 8 μg/ml polybrene (Sigma-Aldrich) and mixed with concentrated virus. The cell-virus mixture was spinoculated 1 h at 600 x g 32 °C followed by 2–4 h incubation at 37°C, after which they were collected and plated on 60% Matrigel overlaid with transduction medium without polybrene. Transduced organoids were selected with 2 rounds of 2 μg/ml of puromycin (Sigma-Aldrich) on day 2 and day 4, after which clones were expanded in maintenance ENR medium. Knockout was confirmed by western blot to check for the expression of deleted gene.

Oligonucleotides used for generation of gRNAs: Esrrg (1) CACCGTCTGTCAAGACGGACCCCTG, AACCAGGGGTCCGTCTTGACAGAC, Esrrg (2) CACCGTGGCGTCGGAAGACCCACCA; AAACCAGGGGTCCGTCTTGACAGAC. Spi-B (1) CACCGAGACTCCTTCTGGGTACTGG, AAACCCAGTACCCAGAAGGAGTCTC; Rank (1) CACCGAAAGCTAGAAGCACACCAG, AAACCTGGTGTGCTTCTAGCTTTC.

### Lentivirus infection for overexpression

RelB, p52, RelA, p50 plasmids were a gift from Hiroshi Ohno’s lab (RCIMS, Kangawa, Japan) and Esrrg cDNA was cloned by Twist bioscience (California, USA). These were cloned into CSII-CMV-MCS-IRES2-Bsd vector which was kindly provided by the RIKEN bioresource center Japan and Hiroyuki Miyoshi. The same protocol for Crispr-Cas9 lentiviral generation and transduction was followed and cells were embedded into Matrigel and incubated for 2-3 days.

### Immunoblotting

Organoids were recovered from Matrigel with Cell Recovery media (Corning). Organoids were washed with PBS and the cells were lysed with 2x Laemmli solution and boiled at 98 degrees. Protein concentrations were measured by Pierce 660nm Protein Assay Reagent and IDCR (ThermoFisher Scientific,22660). Samples were loaded equally in terms of protein concentration into 10% Bis-Tris protein gels (Life Technologies) and blotted on nitrocellulose membranes. Membranes were incubated with primary antibodies: Anti-Esrrg (abcam, ab49129); Anti-Spi-B (Spi-B D4V9S, CST 14337); Anti-H3K27me3 (abcam, ab192985); Anti-H3 (abcam, ab1791), Anti-Rank (MyBioSource, MBS9133424), Anti-GAPDH (abcam, ab8245) at 4 °C overnight, and HRP-conjugated anti-rabbit (Sigma-Aldrich; 1:5000) or anti-mouse (CST; 1:1000) for 1 h at room temperature. Signal was detected using ECL reagent (Amersham 2232).

### Real-time quantitative reverse transcription PCR

Total RNA was prepared using TRIzol (Life Technologies) from intestinal organoids and epithelium isolated from mice. Isolated RNA was transcribed to first strand cDNA using iScript cDNA synthesis Kit (Biorad, 1708891). qPCR amplification was detected using Ssofast evergreen supermixes (Biorad,172-5203). The specific primers used, are listed in the supplementary data.

## Data availability

The GhIP- and Gro-seq data have been deposited in the NCBI Gene Expression Omnibus database (GSE157629).

## Acknowledgements

We would like to thank Rafael Jiménez for providing Sox8-/- mouse transgenic line. This work was supported by the Academy of Finland (no. 310011), Tekes (Business Finland) (no. 658/31/2015), Paediatric Research Foundation, Sigrid Jusélius Foundation, Mary och Georg C. Ehrnrooths Stiftelse, Laboratoriolääketieteen Edistämissäätiö sr. The funding sources played no role in the design or execution of this study or in the analysis and interpretation of the data.

## Author contributions

JJG, KV: Study concept and design. JJG, LMD, VZ, SI, TR, FTAM, HN: experiments and acquisition of data. MO: bioinformatic analyses. JJG, LMD, KV: analysis and interpretation of data. JJG, MO, KV: manuscript drafting. SI, PK, HN, MK: data curation & reviewing the manuscript KV: project administration and funding acquisition. JJG, MO, LMD, VZ, SI, TR, FTAM, HN, PK, MK, KV: critical revision of the manuscript for important intellectual content. All authors approved the final version of the manuscript.

---

**Supplementary table 1**. Gro-seq data for the organoids treated with Rankl and compared to organoids grown in enterocyte differentiation (ENRI) and and stemness (WENRC) conditions.

**Supplementary table 2**. H3K27me3 ChIP-seq data for the organoids treated with Rankl and grown in stemness condition (WENRC)

