## Supplementary figure 1 & 2 for "Polycomb Repressive Complex 2-controlled Essrg regulates intestinal Microfold cell differentiation"

Supplementary Figures

A

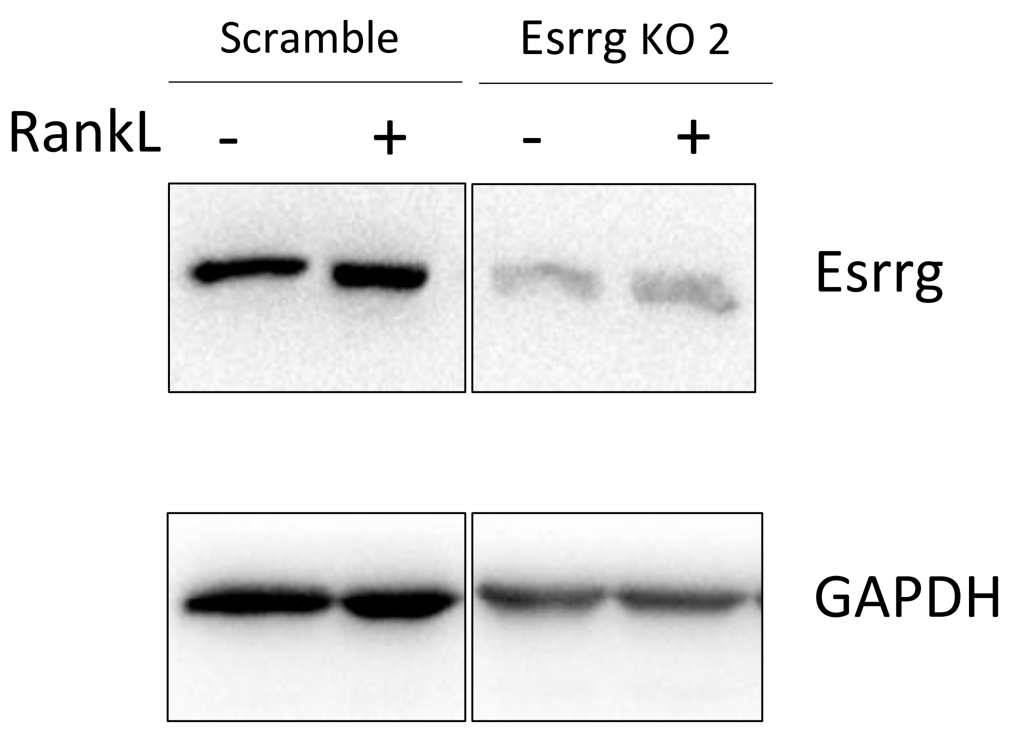


B

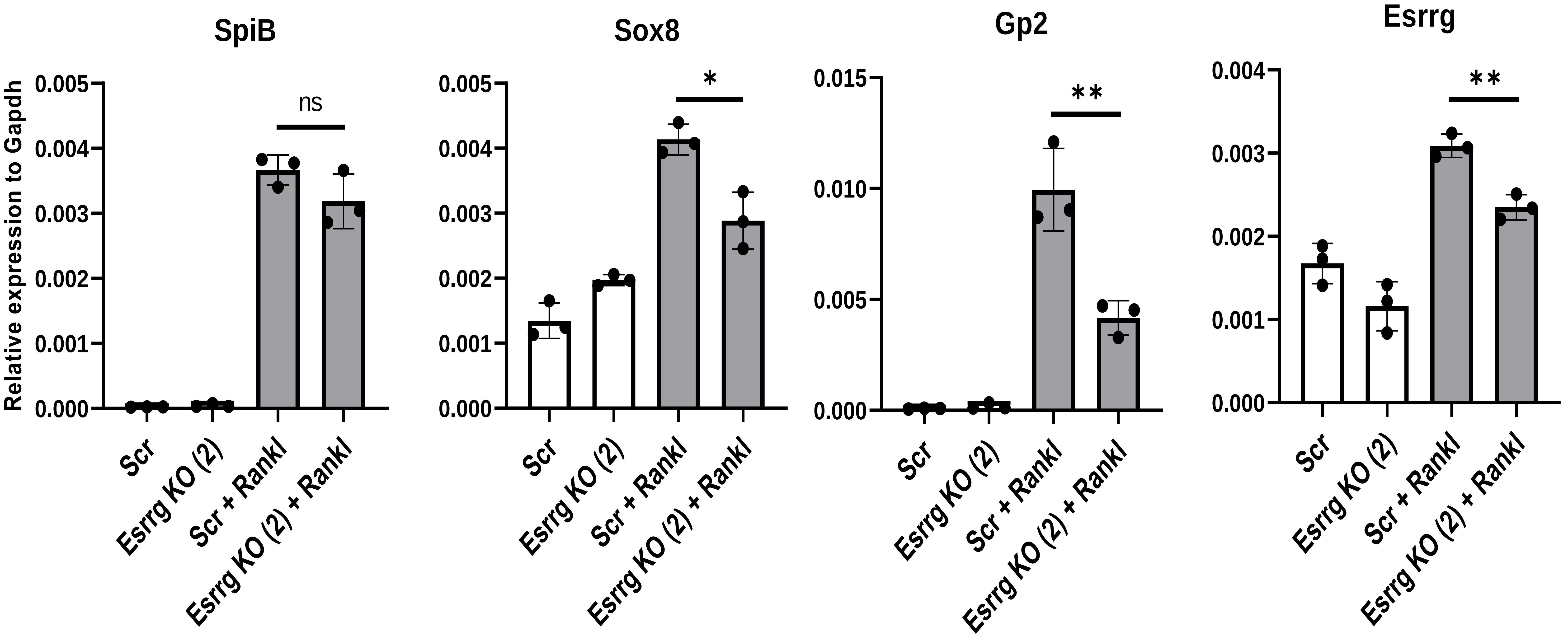


(A)Esrrg protein expression with a second guide RNA targeting Esrrg in Esrrg KO and Scrambled cells generated by CRISPR-Cas9 genome editing in C57BL/6rj intestinal organoids. Organoid lysates were analyzed by Western blot. (B) qPCR analysis of M cell associated transcription factors, Spi-B, Sox8 and GP2 expression in Esrrg KO with guide RNA 2. Results were similar to Esrrg KO generated with guide RNA2; n.s, not significant; *, P< 0,05; **, P < 0.01. unpaired two-tailed Student’s t test; n = 3. Values are presented as the mean ± SD. Data are representative of three independent experiments.

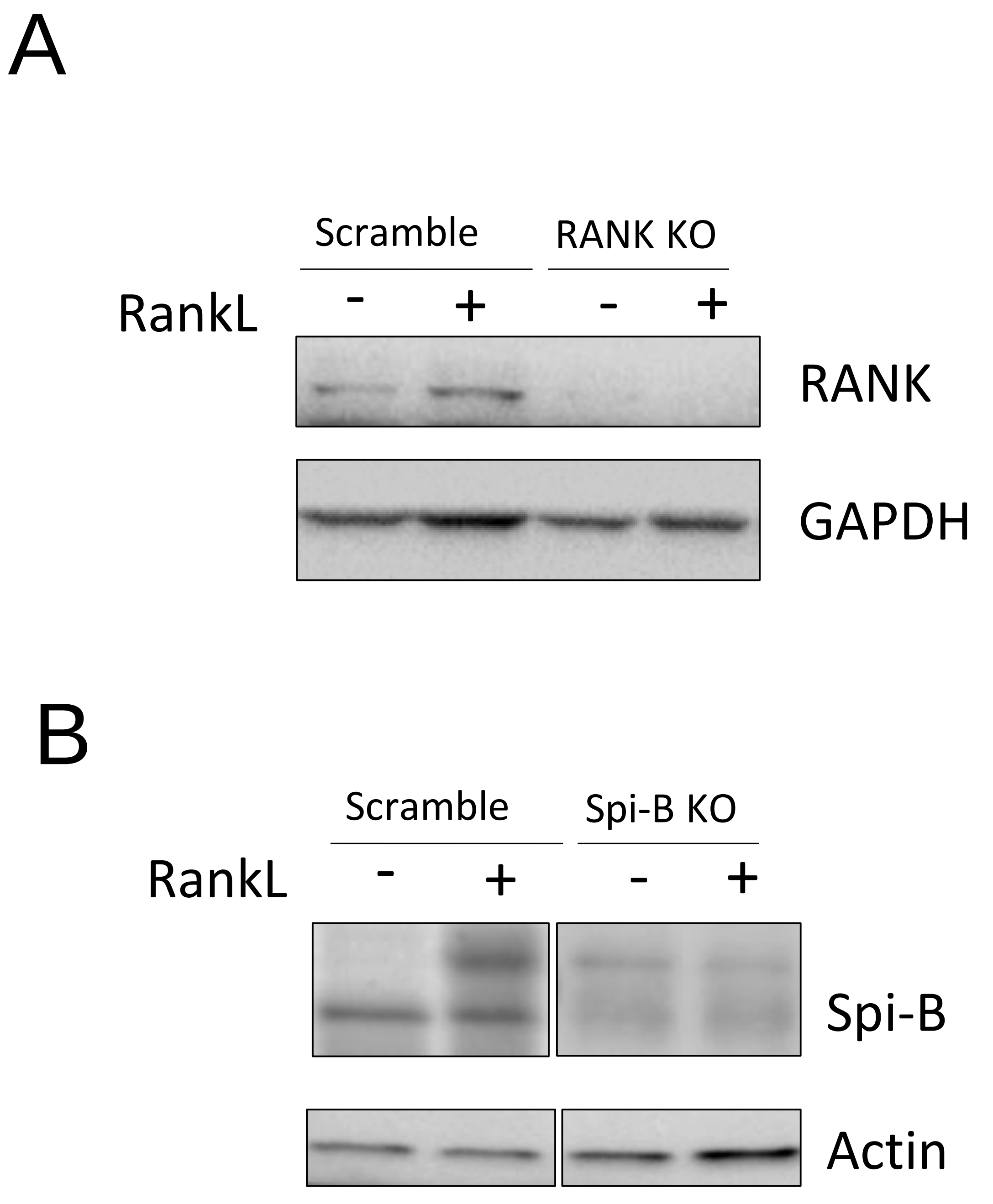


(A) Spi-B KO validation by immunoblot analysis fof SpiB KO and Scrambled intestinal organoids generated by CRISPR-Cas9 genome editing in C57BL/6rj intestinal organoids. These organoids were grown with and without RankL 100ng for 4 days(B) RANK KO validation by immunoblot analysis of RANK KO and Scrambled intestinal organoids generated by CRISPR-Cas9 genome editing in C57BL/6rj intestinal organoids. These organoids were grown with and without RankL 100ng for 4 days

List of oligonucleotide Primers used for RT-PCR

| Oligonucleotide | Sequence (5’ to 3’) |
| --- | --- |
| Gapdh_fwd | TGTGTCCGTCGTGGATCTGA |
| Gapdh_rev | CCTGCTTCACCACCTTCTTGA |
| Suz12_fwd | GATGAGAAAGATCCAGAATGGC |
| Suz12_rev | ATAATTTTCTACAAACAGCATACAGGC |
| Ezh2_fwd | GTCTGATGTGGCAGGCTGG |
| Ezh2_rev | GCCCTTTCGGGTTGCATC |
| Spi-B_fwd | GGAGTCTTCTACGACCTGGACAG |
| Spi-B_rev | GCAGGATCGAAGGCTTCATAGG |
| Sox8_fwd | GGACCAGTACCCGCATCTCC |
| Sox8_rev | TTCTTGTGCTGCACACGGAGC |
| GP2_fwd | GTGTACAAGTTACAGGGTACCCC |
| GP2_rev | GACAAGTAATCTCACAATTCTTGG |
| CCL9_fwd | GCCCAGATCACACATGCAAC |
| CCL9_rev | AGGACAGGCAGCAATCTGAA |
| MarcksL1_fwd | CCCGTGAACGGAACAGATGA |
| MarcksL1_rev | CCCACCCTCCTTCCGATTTC |
| Esrrg_fwd | GTGTCTCAAAGTGGGCATGC |
| Esrrg_rev | GCTGTTCTCAGCATCTATTCTGC |
| Aif1_fwd | GGATTTGCAGGGAGGAAAA |
| Aif1_rev | TGGGATCATCGAGGAATTG |
| CCL20_fwd | TGTACGAGAGGCAACAGTCG |
| CCL20_rev | TCTGCTCTTCCTTGCTTTGG |
| TNFAIP2_fwd | GTGCAGAACCTCTACCCCAATG |
| TNFAIP2_rev | TGGAGAATGTCGATGGCCA |
| 18s rRNA_fwd | GTAACCCGTTGAACCCCATT |
| 18s rRNA_rev | CCATCCAATCGGTAGTAGCG |
